# Haplotype Analysis of the TRB Locus by TCRB Repertoire Sequencing

**DOI:** 10.1101/406157

**Authors:** Timothy J Looney, Dzifa Y Duose, Geoffrey Lowman, Elizabeth Linch, Joud Hajjar, Denise Topacio-Hall, Mingxuan Xu, Jianping Zheng, Anas Alshawa, Coya Tapia, Bettzy Stephen, Linghua Wang, Funda Meric-Bernstam, Lauren Miller, Alexander Glavin, Lifeng Lin, Jing Gong, Jeffrey Conroy, Carl Morrison, Fiona Hyland, Aung Naing

## Abstract

Polymorphism within the T cell receptor beta variable gene (TRBV) has been implicated in autoimmune disease and immuneCrelated adverse events (IRAEs) during immunotherapy. Previous efforts to evaluate TRBV polymorphism by whole genome sequencing (WGS) have been hampered by the repetitive nature of the TCRB locus. We present a novel longCamplicon TCRB repertoire sequencing approach to evaluate TRBV polymorphism from peripheral blood, which we use to identify TRBV allele haplotypes in 81 Caucasians.

## Main

Checkpoint blockade immunotherapy (CPI) can elicit antiCcancer T cell responses mediating durable progression free survival but may also promote T cell destruction of healthy tissue to elicit IRAEs. Efforts to identify germline variants associated with adverse events using WGS or microarrays have yet to reveal markers predictive of adverse events following immunotherapy (Gowen). Identifying such biomarkers could allow for personalized drug selection and dosing to ultimately enable safer and more effective immunotherapy, particularly in light of the increasing use of combination CPI regimens having a significant incidence of severe adverse events (Larkin). Despite previous efforts, three lines of reasoning support the notion that germline encoded TRBV polymorphism could be a key determinant of adverse events during CPI. First, the TCR locus is repetitive and structurally complex, impeding the measurement of variation by traditional short read WGS or microarray based methods (Watson); second, single amino acid substitutions within the framework or CDR 1 and 2 regions of the rearranged TCRB chain are know to significantly alter TCR affinity for HLA (Gras, Robbins 2008, Robbins 2011); and third, adverse events during immunotherapy may manifest as acute versions of chronic autoimmune diseases (i.e. fulminant type 1 diabetes) that have been separately linked to TRBV polymorphism (Pierce, Hockertz, McDermott, Maksymowych, Gaudi, Hughes, Okamoto). To circumvent the challenge in measuring TRBV polymorphism by WGS, we developed a method for detection of TRBV polymorphism by nextCgeneration sequencing (NGS) of rearranged TCRB chains from peripheral blood leukocytes (PBL). To our knowledge this represents the first NGSCbased method to permit haplotypeClevel resolution of the TRB locus.

Our workflow begins with the cDNA derived from PBL total RNA, which is used as input for multiplex PCR via framework 1 (FR1) and constant gene primers to generate ~330bp TCRB amplicons spanning the three beta chain CDR regions.

This extended amplicon enables detection of polymorphism within the germlineC encoded framework and CDR1 and 2 regions in addition to the highly variable CDR3 region (Figures 1A and B). To minimize primerCprimer interaction and amplification bias, we generated primer sequences in accordance with AmpliSeq design principles (Merriman) to cover known population variants and ensure robust and reproducible amplification of all alleles. Following amplification, primers are digested and resultant amplicons ligated to oligomer adaptors for bidirectional NGS. We choose to employ Ion Torrent sequencing via the S5 530 chip, which is favorable for this application owing to the long read lengths (up to 600bp) and low substitution error rate (Merriman); our estimates from sequencing of bacterial genomes indicate that the substitution error rate may be up to a magnitude lower than that of commonlyCused Illumina machines (Supplemental Figures 1A and B). As discussed in depth below, substitution errors pose a key challenge to immune repertoire analysis given that they may mimic the natural variation in the repertoire.

**Figure 1.**
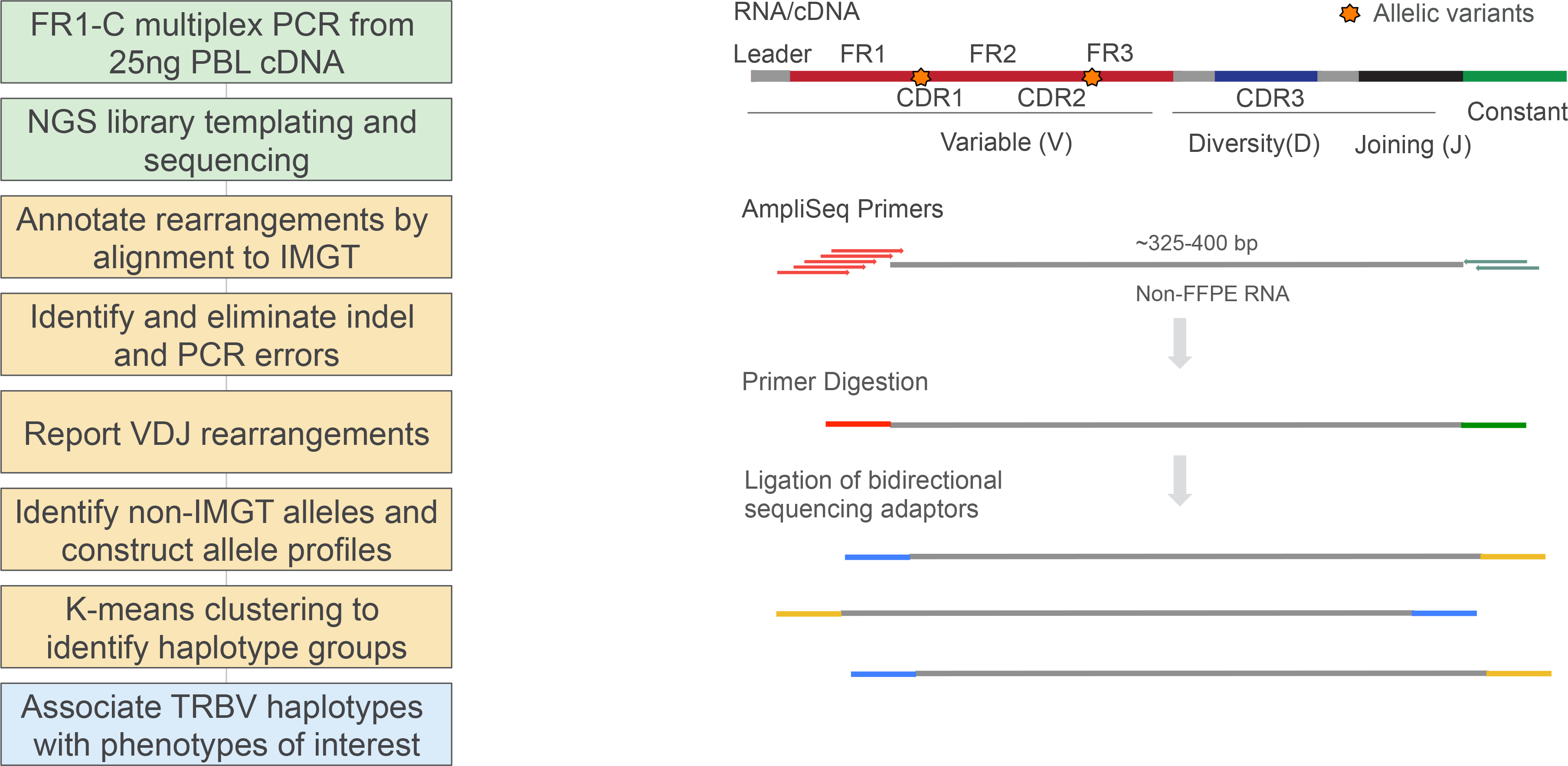
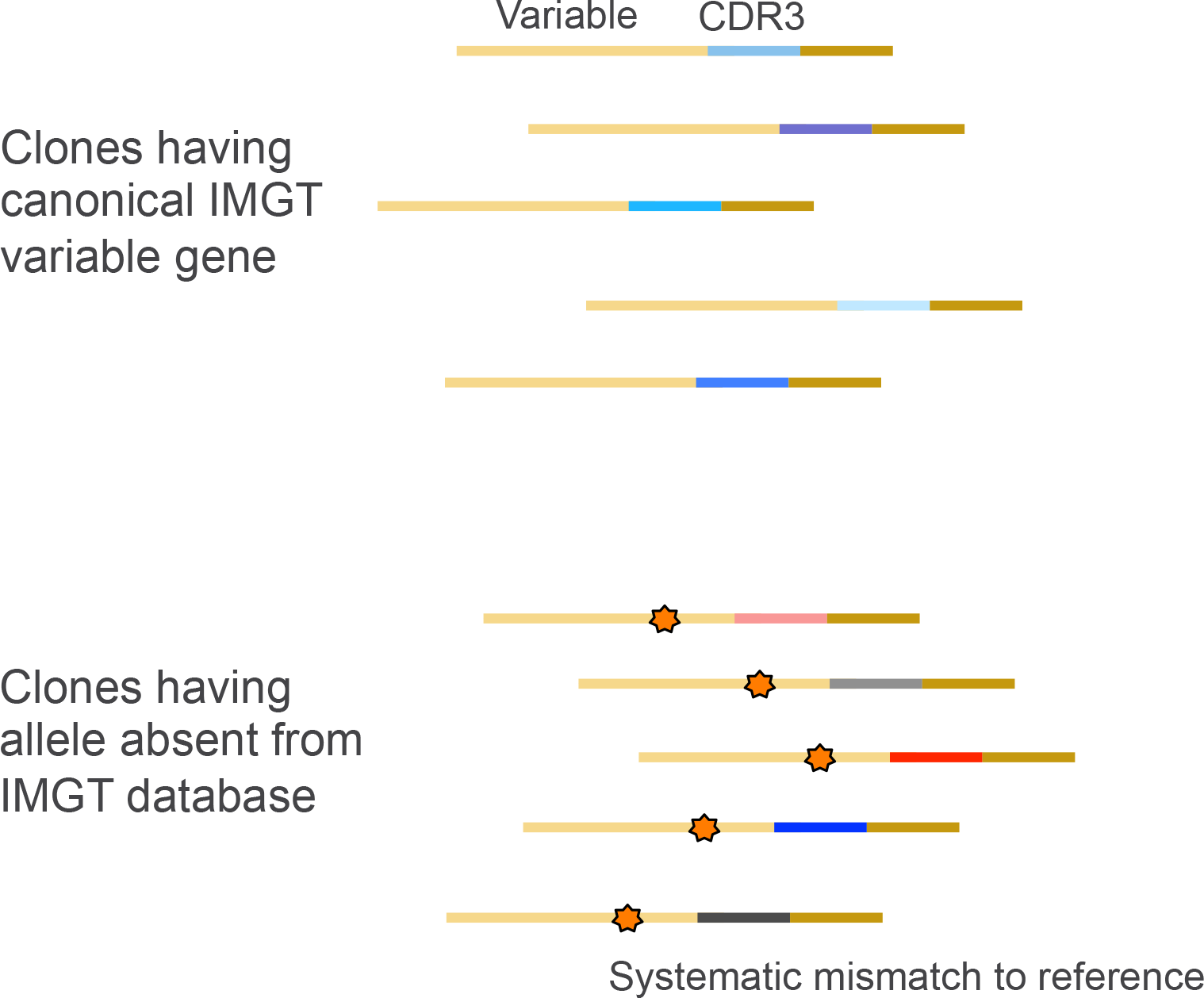
**(A)** Overview of workflow. The reagents and informatics methodology have been incorporated into the publicly available Oncomine TCR BetaCLR assay and accompanying Ion Reporter analysis software. **(B)** Method for library preparation and sequencing. Our strategy utilizes multiplex PCR via framework 1 and constant gene primers followed by bidirectional NGS of amplicon libraries. **(C)** Strategy for identification of nonCIMGT variable gene alleles. Bone fide novel alleles will present as systematic mismatches to IMGT across a plurality of clones, each possessing a distinct CDR3 nucleotide sequence.

Following sequencing our workflow maps sequence reads to variable, diversity and joining genes from the IMGT database and eliminates reads representing offCtarget products or those that do not span the entire FR1CConstant region of the amplicon. The Ion Torrent instruments have an error mode dominated by homopolymer errors manifesting as a single base expansion or reduction of an existing homopolymer tract. To identify sequences with indel errors, we translate VDJ sequences to protein space and evaluate the productivity of the rearrangement, i.e. whether the V and J gene are in frame and there are no premature stop codons. Indel errors manifest as frameshift mutations that make rearrangements appear unproductive. Owing to nonsense mediated decay, T cell allelic exclusion and thymic selection, it is expected that unproductive rearrangements should be exceedingly rare in RNA and thus one may infer that unproductive sequences contain a sequencing error. In some cases, an indel error may occur within the variable gene portion of the sequence such that alignment to the variable gene IMGT reference indicates a gap within the homopolymer tract containing the indel error. ReC evaluation of the initial base calling at such position often leads to correction of the error to produce an ostensibly sequencing errorCfree read representing a productive rearrangement. After indel error correction we eliminate PCRCderived errors by evaluating edit distances between VDJ rearrangements, taking into account the frequency of the rearrangement and whether the rearrangement is supported by forward strand and reverse strand reads. Finally, the pipeline reports the sequence and frequency of IMGTCannotated VDJ rearrangements detected in the sample.

We next use the set of annotated VDJ rearrangements to determine the variable gene allele profile of an individual. Though the IMGT database is considered to be gold standard, evidence (Yu) suggests that it is incomplete. To identify instances where an individual possesses a nonCIMGT (putatively novel) variable gene allele, we leverage the fact that clones utilizing a nonCIMGT allele will present as having a systematic mismatch to the IMGT database. Given that each clone is readily distinguishable from one another in sequence space owing to the diversity of the CDR3 region, the number of clones having a particular systematic mismatch is indicative of the minimum number of unique template molecules supporting aputative nonCIMGT allele. Bone fide novel alleles will be found on a plurality of clones, each possessing a distinct CDR3 nucleotide sequence, while mismatches owing to random PCR error or sequencing error will not be found on multiple clones within a repertoire (Figure 1C). To report an allele for downstream analysis, either a putative novel allele or canonical IMGT allele, we require that the allele be present on a minimum of 5 clones and make up at least 5% of the sequences obtained for that variable gene. We allow for up to two alleles of a particular variable gene to be detected in a single sample. In the hypothetical case where more than two potential alleles are detected for a particular variable gene, only the two alleles having the greatest clone support are reported.

Owing to genetic linkage and population structure, we expect to observe setsof coCinherited variable gene alleles (i.e. allele haplotypes) within human populations. To identify such haplotypes we applied our method to obtain TRBV allele profiles for 81 Caucasians, then combined the profiles into a matrix such that each row of the matrix represents a different individual and each column represents a different variable gene allele, where ‘1’ indicates presence of allele and ‘0’ indicates allele absence (Supplemental Table 1). Our pipeline identified 37 nonC IMGT variable gene alleles in the sample set, of which 19 are found in less stringent databases such as Lym1k (Yu) or the NCBI NR archive. The remaining 18 variable gene alleles appear novel to literature; sequences of these alleles are provided in Supplemental Table 2. We next performed principal component analysis of the matrix of variable gene allele profiles to extract the two largest components contributing to differences in allelic representation among the sample set. KCmeans clustering of principal components revealed 6 major allele profiles within the dataset, which we term haplotype groups (Figures 2A and B). The optimal number of clusters was determined using the “elbow” method and plotting the within cluster sum of squares over cluster centers from 1 to 15 (Supplemental Figure 2).

**Figure 2.**
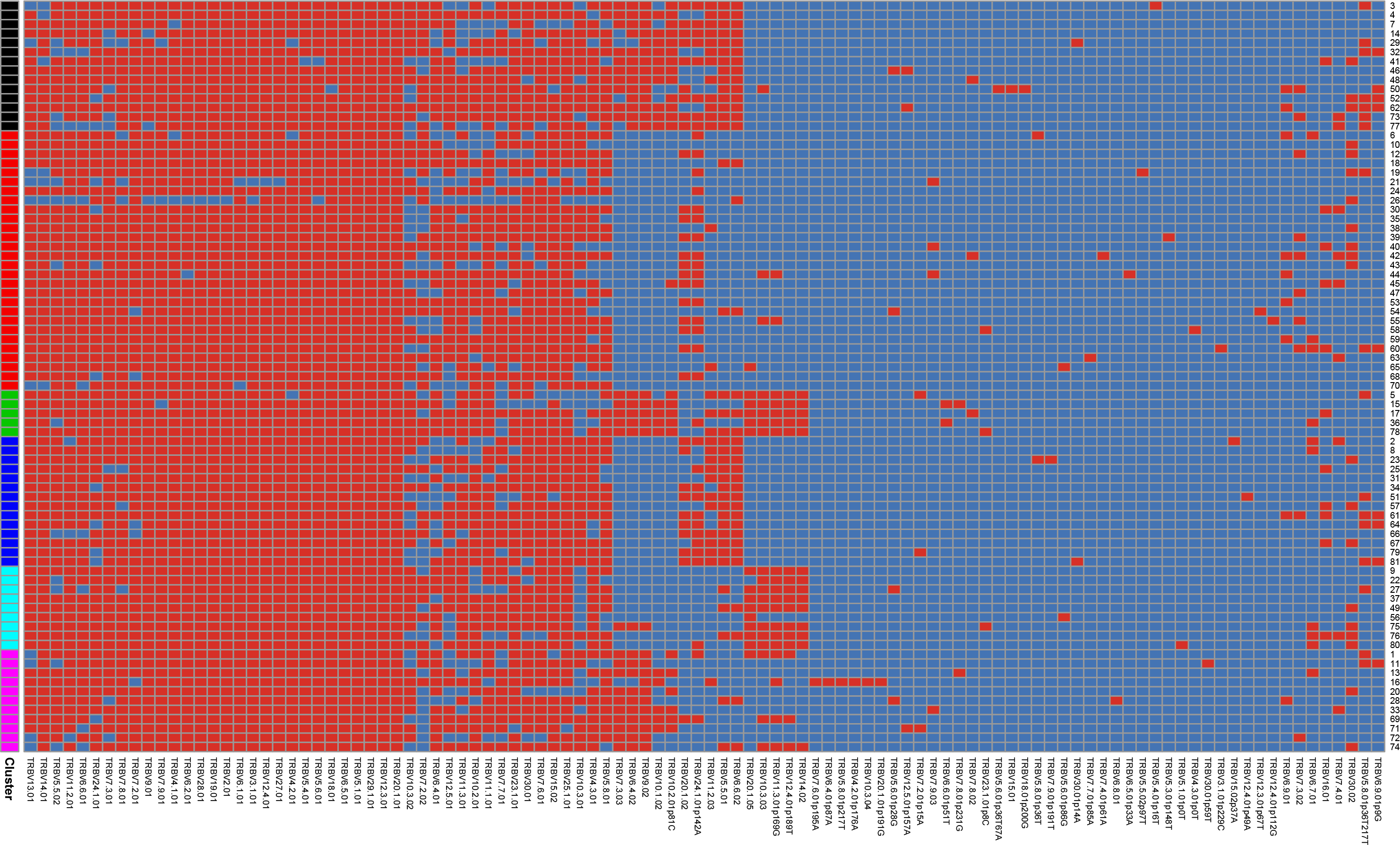
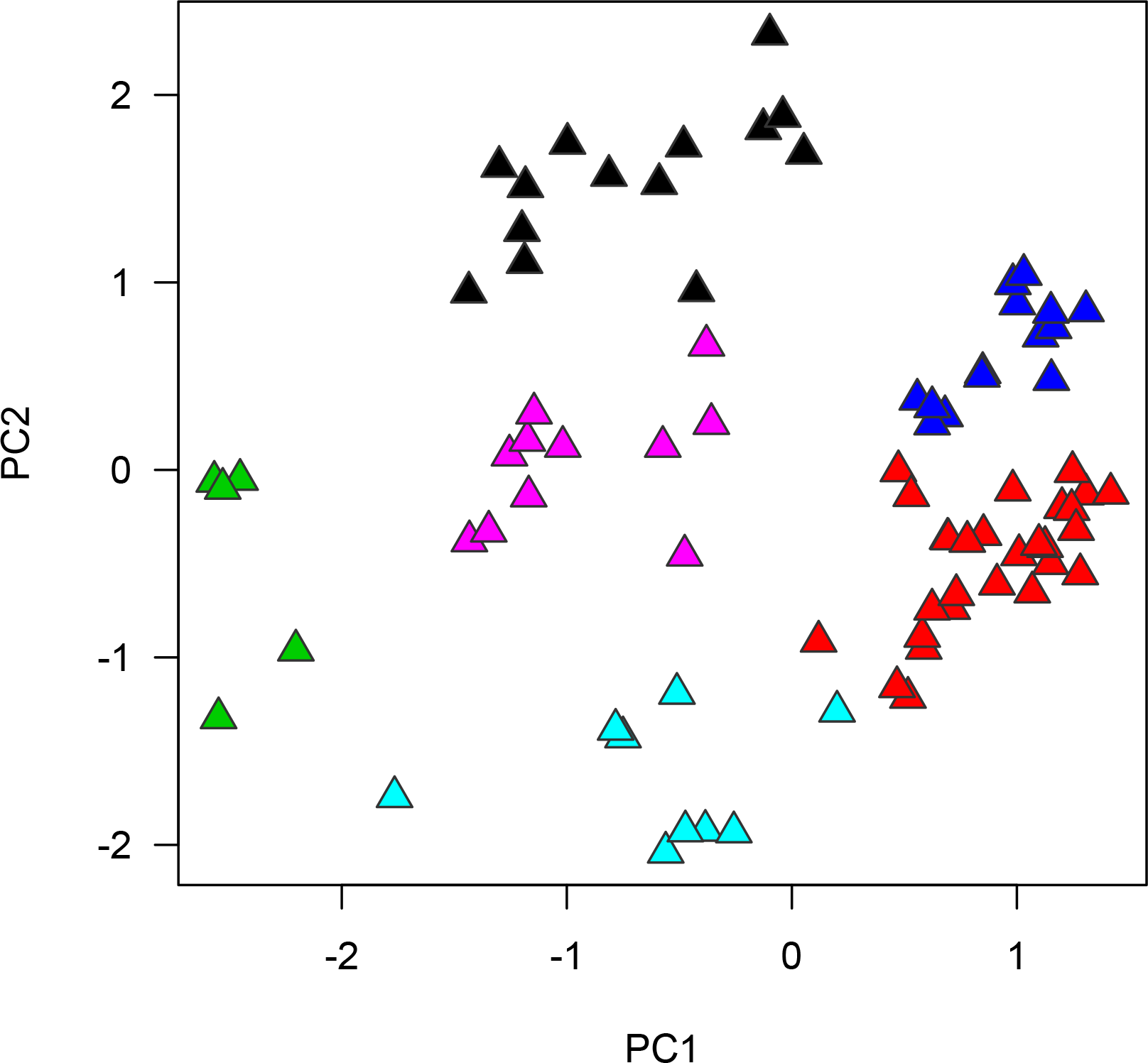
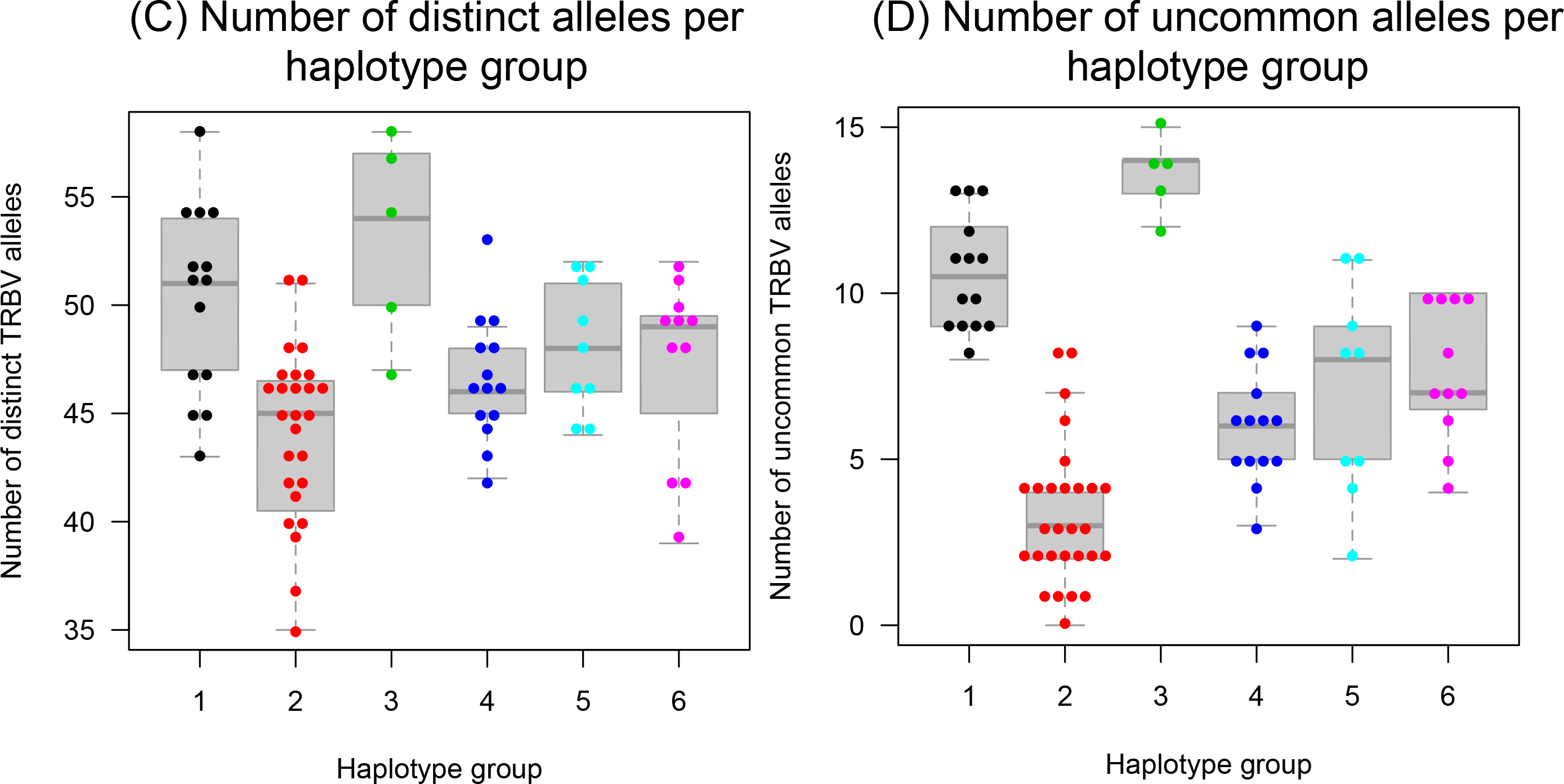
**(A)** Heatmap of TRBV allele profiles for 81 Caucasians. TCRB repertoires were used to construct variable gene allele profiles for each individual. The sets of alleles detected for each individual are displayed in heatmap form, where each row represents a different individual and each column a different variable gene allele. Red tiles indicate that an allele was detected in an individual while blue tile indicate allele absence. Columns are arranged via hierarchical clustering, while rows are arranged according to haplotype group classification produced by kCmeans clustering. IMGT allele names are displayed along the XCaxis; alleles having a lowercase ‘p’ in the name correspond to putative novel alleles absent from the IMGT database. To the left, Cluster column indicates the haplotype group classification. **(B)** Principal component analysis of allele profiles. Samples are displayed according to the two largest principal components derived from analysis of the TRBV allele profile matrix. Samples are colored according to haplotype group classification identified via kCmeans analysis. **(C)** Number of distinct alleles detected across haplotype groups. The number of distinct alleles detected in each individual was calculated then displayed according to haplotype group. **(D)** Number of uncommon alleles detected across haplotype groups. Uncommon alleles are defined as those found in less than 50% of this Caucasian cohort.

We finally asked whether there were allele profile features that distinguish the haplotype groups from one another. We found that haplotype group 2 members (Figure 2B, red) have fewer distinct alleles and fewer uncommon alleles (defined as those present in <50% of the sample set) than members of other haplotype groups, indicating that members of this group tend to be homozygous for a common Caucasian allele haplotype, while members of other groups have elevated TRBV allele heterozygosity and carry TRBV haplotypes that are less common in Caucasians (Figures 2C and D; p=1.7EC4 and 3.6EC13 for number of distinct alleles and uncommon alleles respectively, student’s tCtest). This finding may be significant given proposals regarding the existence of balanced functional polymorphisms within antigen receptor genes, which posit that lower frequency alleles may have a greater tendency towards autoreactivity than the most common alleles in human populations (Dean). Extending on this notion, haplotype group 2 members may be at a lower risk of autoimmune disease and IRAEs than members of other haplotype groups.

In summary, we have developed a costCefficient and rapid method for routine analysis of germlineCencoded polymorphism within the TRB locus to enable highC resolution studies of genetic variation and its potential link to IRAEs, chronic autoimmune disease and response to infectious disease. The reagents and informatics methodology described herein have been incorporated into the publicly available Oncomine TCR BetaCLR assay and accompanying Ion Reporter analysis software. Current and future research will clarify the potential role of TRBV polymorphism in IRAEs and chronic autoimmune disease.

---

**Supplemental Figure 1.**
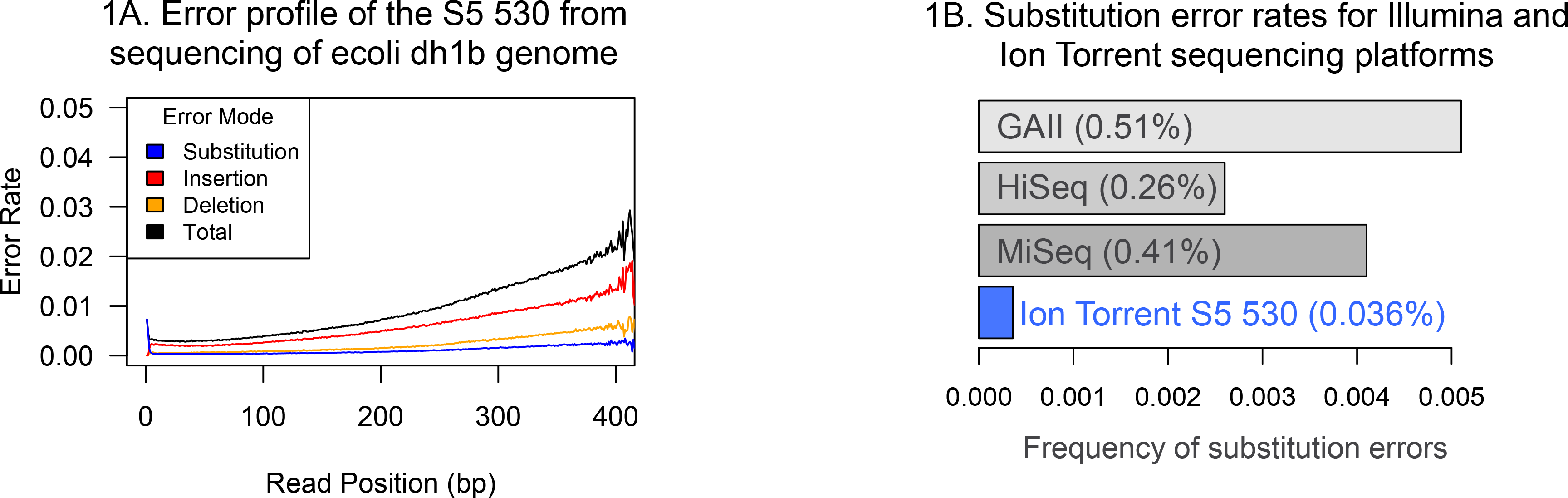
Substitution Error Rates for Ion Torrent and Illumina Machines.

**(A)** The average substitution error rate for the S5 530 chip was calculated by sequencing of the ecoli dh1b genome and mapping sequence reads to reference via the Torrent Accuracy plugin for Torrent Suite. **(B)** Comparison of substitution error rates across three Illumina sequencers and the Ion Torrent S5 530 chip. To account for differences in read lengths between the Illumina and Ion Torrent platforms, the error rate was calculated as the average over the bases 10C 100 of the read. The first 10 base pairs were excluded given that alignment scores cannot accurately distinguish indel errors from substitution errors at the read ends. Error rates for the Illumina platform are taken from Schirmer et al, 2016.

**Supplemental Figure 2.**
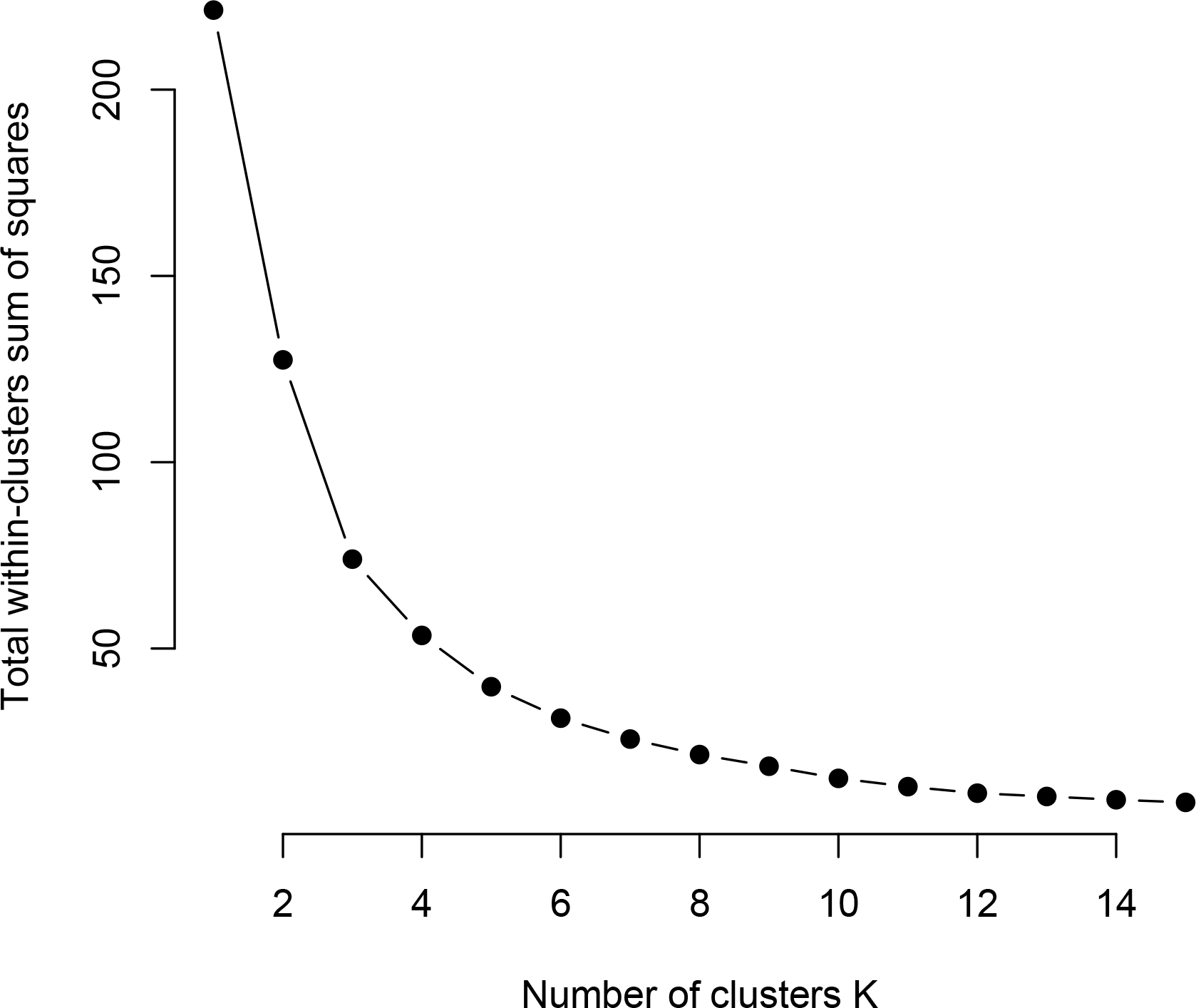
Within Sum of Squares Analysis of K-Means Clustering.

Within cluster sum of squares following subdivision of data into 2 to 15 clusters via kCmeans analysis of sample set.

**Supplemental Table 1.** TRBV allele profiles for samples. Allele profiles were generated as described in methods. Reads column indicates the number of reported reads from Ion Reporter, defined as reads coding for productive rearrangements and corresponding to VDJ sequences that have been determined to be sequencing and PCR error free. PC1 and PC2 columns indicate the loadings from the two largest principal components. Evenness column indicates the normalized Shannon entropy of clone sizes, calculated from the frequencies of detected clones.

**Supplemental Table 2**. Summary of novel alleles detected in dataset. Table indicates sequences of putatively novel alleles, whether an allele is found in the Lym1k database or the NCBICNR database, whether an allele is synonymous in amino acid space with an IMGT allele, and the location of the variant residue within the variable gene, where applicable.

## References

1. Dean, M et al. Balanced polymorphism selected by genetic versus infectious human disease. Annu Rev Genomics Hum Genet (2002). 3:263–292.

2. Gaudi, C et al. AntiCPD1 Pembrolizumab can induce exceptional fulminant type 1 diabetes. Diabetes care (2015). 38:e182C183.

3. Gowen, M et al. Baseline antibody profiles predict toxicity in melanoma patients treated with immune checkpoint inhibitors. J Transl Med (2018). 16:82

4. Gras, S et al. Allelic polymorphism in the T cell repertoire and its impact on immune responses. J Exp Med (2010). 207:155–1567

5. Hockertz, M et al. Susceptibility to relapsingCprogressive multiple sclerosis is associated with inheritance of genes linked to the variable regin of the TCR beta locus: use of affected familyCbased controls. Am. J. Hum. Genet (1998). 62:373–385.

6. Hughes, J et al. Precipitation of autoimmune diabetes with anti PDC1 Immunotherapy. Diabetes Care (2015). 38:e55–57

7. Larkin, J et al. Combined Nivolumab and Ipilimumab or monotherapy in untreated melanoma. NEJM (2015). 373:23–34.

8. Maksymowych, W et al. Polymorphism oin a TCcell receptor variable gene is associated with susceptibility to a juvenile rheumatoid arthritis subset. Immunogenetics (1992). 35:257–262.

9. McDermott, M et al. The role of T cell receptor beta chain genes in susceptibility to rheumatoid arthritis. Arth Rheum (1995). 38:91–95.

10. Merriman, B et al. Progress in Ion Torrent semiconductor chip based sequencing. Electrophoresis (2012). 33:3397–3417.

11. Okamoto, M et al. Fulminant type 1 diabetes mellitus with antiCprogrammed cell deathC1 therapy. J Diabetes Investig (2016). 7:915–918.

12. Pierce, B et al. The missing heritability in T1D and potential new targets for prevention. J Diabetes Res (2013). 2013:737485.

13. Robbins, P et al. Single and dual amino acid substitutions in TCR CDRs can enhance antigenCspecific T cell functions. J Immunol (2008). 180:6116–6131.

14. Robbins, P et al. Tumor regression in patients with metastatic synovial cell sarcoma and melanoma using genetically engineered lymphocytes reactive with NYCESOC1. J Clin Oncol (2011). 29:917–924.

15. Schirmer et al. Illumina error profiles: resolving fineCscale variation in metagenomic sequencing data. BMC Bioinformatics (2016). 17:125.

16. Stucci, S et al. ImmuneCrelated adverse events during anticancer immunotherapy: pathogenesis and management (review). Oncol Letters (2017). 14:5671:5680.

17. Watson, C et al. Comment on ‘A database of human immune receptor alleles recovered from population sequencing data’. J Immunol (2017). 198:3371–3373.

18. Yu, Y et al. A database of human immune receptor alleles recovered from population sequencing data. J Immunol (2017). 198:2202–2210.

